# Coping with stress: morphological changes in response to environmental stimuli in a fungal plant pathogen

**DOI:** 10.1101/372078

**Authors:** Carolina Sardinha Francisco, Xin Ma, Maria Manuela Zwyssig, Bruce A. McDonald, Javier Palma-Guerrero

## Abstract

During their life cycles, pathogens have to adapt to many biotic and abiotic environmental stresses to maximize their overall fitness. Morphological transitions are one of the least understood of the many strategies employed by fungal plant pathogens to adapt to constantly changing environments, even though different morphotypes may play important biological roles. We characterized the responses of the wheat pathogen *Zymoseptoria tritici* to a series of environmental stresses in order to understand the effects of changing environments on fungal morphology and adaptation. We found that all tested stresses induced morphological changes, but different responses were found among four different strains. A transcription analysis showed that morphogenesis and virulence factors are co-regulated. We discovered that *Z. tritici* forms chlamydospores and demonstrated that these structures are better able to survive extreme cold, heat and drought than other cell types. We also show that blastospores (the “yeast-like” form of the pathogen typically found only in laboratory conditions) can form from germinated pycnidiospores on the surface of wheat leaves, suggesting that this morphotype can play an important role in the natural history of *Z. tritici*. Our findings illustrate how changing environmental conditions can affect cellular morphology and lead to the formation of new morphotypes, with each morphotype having a potential impact on both pathogen survival and disease epidemiology.

## Introduction

Fungi occupy a wide range of niches that can impose different environmental constraints on their growth, reproduction and survival. In response, fungi evolved the ability to detect and respond to different environmental stimuli to maximize their fitness [1]. One of the strategies that fungi employ to cope with diverse biotic and abiotic stresses is to change their morphology [2–4]. For fungal pathogens, different morphotypes can play different roles during the host-pathogen interaction to optimize overall pathogen fitness. For example, hyphae can be the most appropriate morphology to cross physical barriers, colonize host tissue [2,5], and escape harmful environments generated by host defences [2,6] while yeast cells (or “yeast-like” cells) may be a better morphology to enable dispersal among host niches or between hosts and increase the pathogen population size [2,7]. The majority of fungal morphotype transitions occur between hyphal and yeast growth forms. Fungi with this ability are called dimorphic. Usually, they grow as saprophytic molds that live on dead organic matter and many are facultative human pathogens, though dimorphism is also found in plant pathogens [2,8,9]. The morphological transition can be co-regulated with the expression of genes encoding virulence [2–5,7,10,11]. The induction of virulence genes is usually associated with a specific morphotype. For example, certain virulence genes are expressed only during the yeast-phase of growth in *Histoplasma capsulatum*, while other virulence genes are expressed only during hyphal growth in *Candida albicans* [5,6,12].

Some fungi can produce additional morphologies, including pseudohyphae and chlamydospores. Fungi that can produce more than two morphotypes are called pleomorphic. Pseudohyphae are distinguished from true hyphae by constrictions that form at the septal junctions, leading to a loss in cytoplasmic continuity between mother and daughter cells [13]. Though little is known about the ecological significance of pseudohyphae and their role during host infection, it was suggested that this morphotype facilitates foraging for scarce nutrients and increases motility in the host [7,14]. Chlamydospores are spherical, thick-walled cells formed on the tips of pseudohyphae or on distal parts of differentiated hyphae called suspensor cells [15]. In contrast to pseudohyphae, chlamydospores are well-characterized and known to operate as long-term survival structures that can resist many environmental stresses [15–19]. Pleomorphism has been observed in fungal pathogens of humans [15,17,20] and plants [18,21,22].

*Zymoseptoria tritici* is the most damaging pathogen of wheat in Europe [23,24] and an important pathogen of wheat worldwide. *Z. tritici* is considered a dimorphic fungus that grows either as blastospores (the “yeast-like” form) or as filamentous hyphae depending on the culture conditions. Though the role of blastospores in nature was unclear until now, hyphae are essential for penetrating wheat leaves through stomata [25,26]. The stimuli that induce morphological changes and the cell signaling pathways involved in morphological transitions remain largely unknown in this pathogen. Nutrient-rich media and temperatures ranging from 15°C to 18°C tend to induce blastosporulation (blastospore replication by budding) while nutrient-limited media and high temperatures (from 25°C to 28°C) tend to promote the blastospore-to-hyphae transition in the *Z. tritici* reference strain IPO323 [27,28]. Some genes affecting morphogenesis in *Z. tritici* were identified and functionally characterized. Most of these belonged to the mitogen-activated protein kinase (MAPK) or cAMP-dependent signaling pathways involved in extracellular signal transduction, regulating many cellular processes, development and virulence [27–37]. However, the transcriptome signatures associated with specific morphotypes were not reported until now.

In this study we aimed to characterize how environmental changes affect morphology and adaptation in a fungal plant pathogen. We found that *Z. tritici* changed its morphology in response to all tested stresses, though distinct responses were observed for four different strains sampled from the same geographical population. Transcriptional analyses showed a co-regulation of mycelial growth with expression of virulence factors. We discovered that *Z. tritici* forms chlamydospores both *in vitro* and *in planta*. We also found that pseudohyphae will form in some environments. We showed that blastospores can form by budding from pycnidiospores inoculated onto the surface of wheat leaves, demonstrating that blastospores can be produced under natural conditions. Based on these findings, we propose that morphological transitions in *Z. tritici* are a stress response that allows the fungus to optimize fitness in a changing environment.

## Results

### High temperature induces different morphological responses among *Z. tritici* strains

We exposed four different *Z. tritici* strains growing in yeast sucrose broth (YSB) at 27°C, a highly stressful temperature for *Z. tritici*. Three of the strains (1E4, 3D1, and 3D7) differentiated to filamentous growth at 27°C (Fig 1) as previously reported for the reference strain IPO323 [27,28]. The timing of the hyphal transition ranged from 8 hours after incubation (hai) for 1E4, to 24 hai for 3D1 and 3D7 (Figs 1B and 1C). Branching and hyphal elongation were frequently observed at 48 hai (Figs 1D-1F). Surprisingly, high temperature promoted the formation of two new morphologies, with appearances matching pseudohyphae and chlamydospores, in the 1A5 strain (Figs 1E and 1F). However, no hyphal growth was observed for this strain. Interestingly, chlamydospore-like cells were found for all strains at 144 hai (Fig 1F). Cellular differentiation from blastospores to pseudohyphae began in 1A5 at 24 hai, resulting in a series of conjoined elongated cells, characteristic of pseudohyphae [13] (Fig 1C). We also observed several swollen cells at 24 hai that we believe represent the initial differentiation from blastospores into chlamydospore-like cells (Fig 1C), but the complete differentiation occurred between 48 and 96 hai (Figs 1D and 1E). Chlamydospore-like cells were observed at the ends of pseudohyphae or distal parts of filamentous hyphae (Fig 1F). Chlamydospore-like cells that were detached from suspensor cells were also found (data not shown). Blastospores incubated in YSB at 18°C continued to replicate only as blastospores via blastosporulation and were used as controls (Fig 1A).

**Fig 1.**
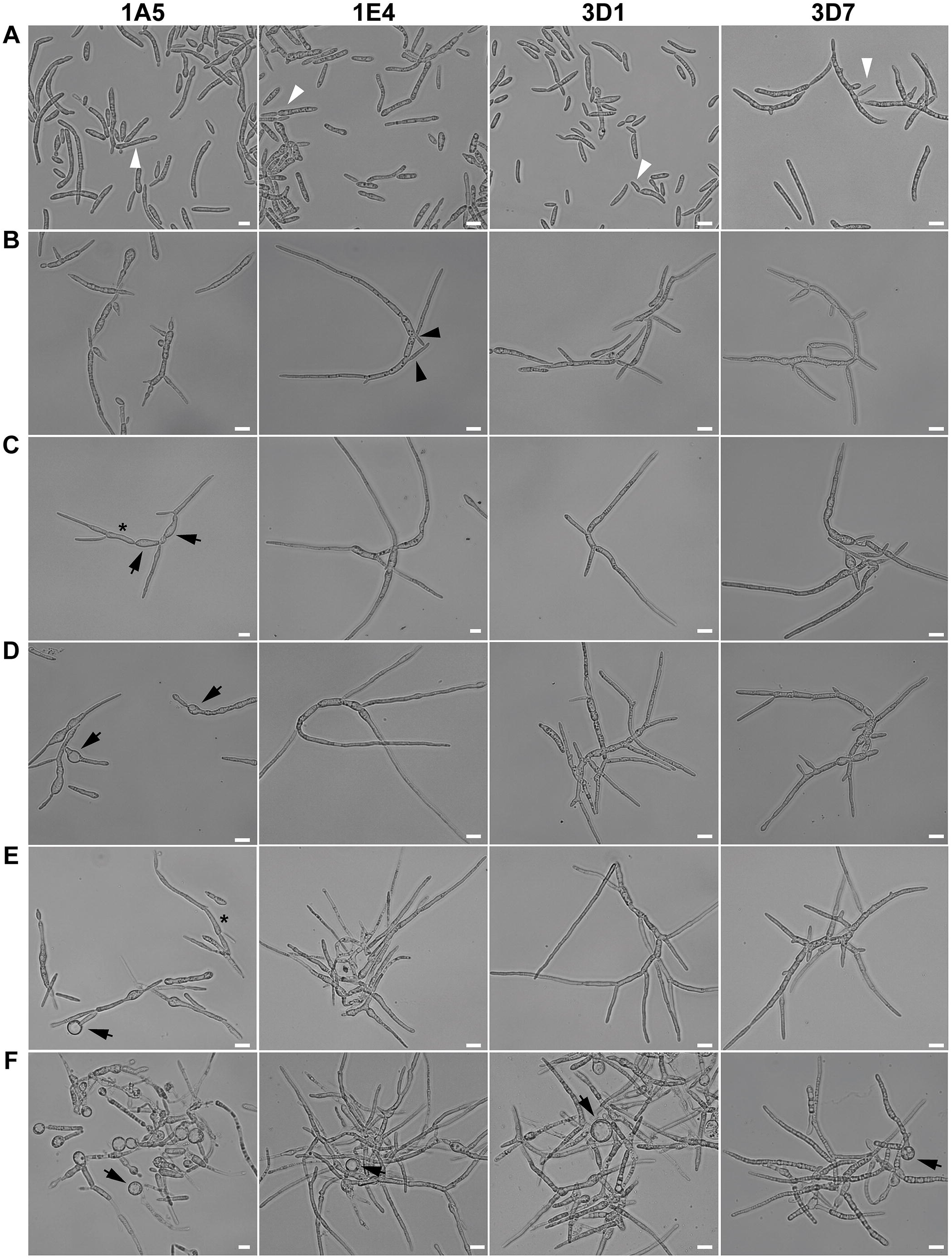
Effect of temperature on the cell morphology of four *Zymoseptoria tritici* strains. (A) Blastospores incubated on nutrient-rich YSB medium at 18°C (the control environment) multiply by budding as blastospores (white triangles). (B) The effect of high temperatures on *Z. tritici* morphology was observed by exposing the strains to 27°C. At 8 hours after incubation (hai) at the high temperature, blastospores of the 1E4 strain were faster to respond to the new environment by switching to filamentous growth. Black triangles point to hyphal branches. (C) At 24 hai, blastospores of the 3D1 and 3D7 strains began growing as hyphae. The 1A5 strain produced swollen cells (black arrows) and pseudohyphae (asterisks), without any evidence of hyphal differentiation. (D) At 48 hai, hyphal branches were abundant for the 1E4, 3D1, and 3D7 strains. The 1A5 strain continued to produce only swollen structures (black arrows). (E) At 96 hai, filamentous growth was observed for the 1E4, 3D1, and 3D7 strains. For the 1A5 strain, high temperature promoted the formation of two new morphotypes: pseudohyphae (asterisks) and chlamydospore-like cells (black arrows). (F) After 144 hours of incubation, the formation of chlamydospore-like cells was observed mainly at the tips of pseudohyphae for the 1A5 strain and at the distal ends of mycelial filaments for the remaining strains (1E4, 3D1, and 3D7) (black arrows). Bars represent 10 μm.

### A nutrient-limited environment promotes filamentous growth

Carbon- or nitrogen-depleted medium promoted hyphal growth in all four strains (Fig 2; S1 Fig). In a carbon-depleted environment (Minimal medium (MM) without sucrose), cellular differentiation occurred after 24 hours of incubation (Fig 2). At 24 hai, hyphal formation was observed with the first hyphal branch formed terminally (1A5 and 3D1) or laterally (1E4 and 3D7) (Fig 2B). Hyphal elongation and branching continued at 48 and 72 hai (data not shown). By 144 hai mycelial growth was predominant, with frequent anastomosis events among hyphae (Fig 2C). In MM with sucrose, all strains grew as a mixture of blastospores and hyphae, with blastospores budding from filamentous hyphae, showing that the morphological transition is bidirectional (S1C Fig). Blastospores incubated in YSB continued to replicate only as blastospores via blastosporulation (Fig 2A).

**Fig 2.**
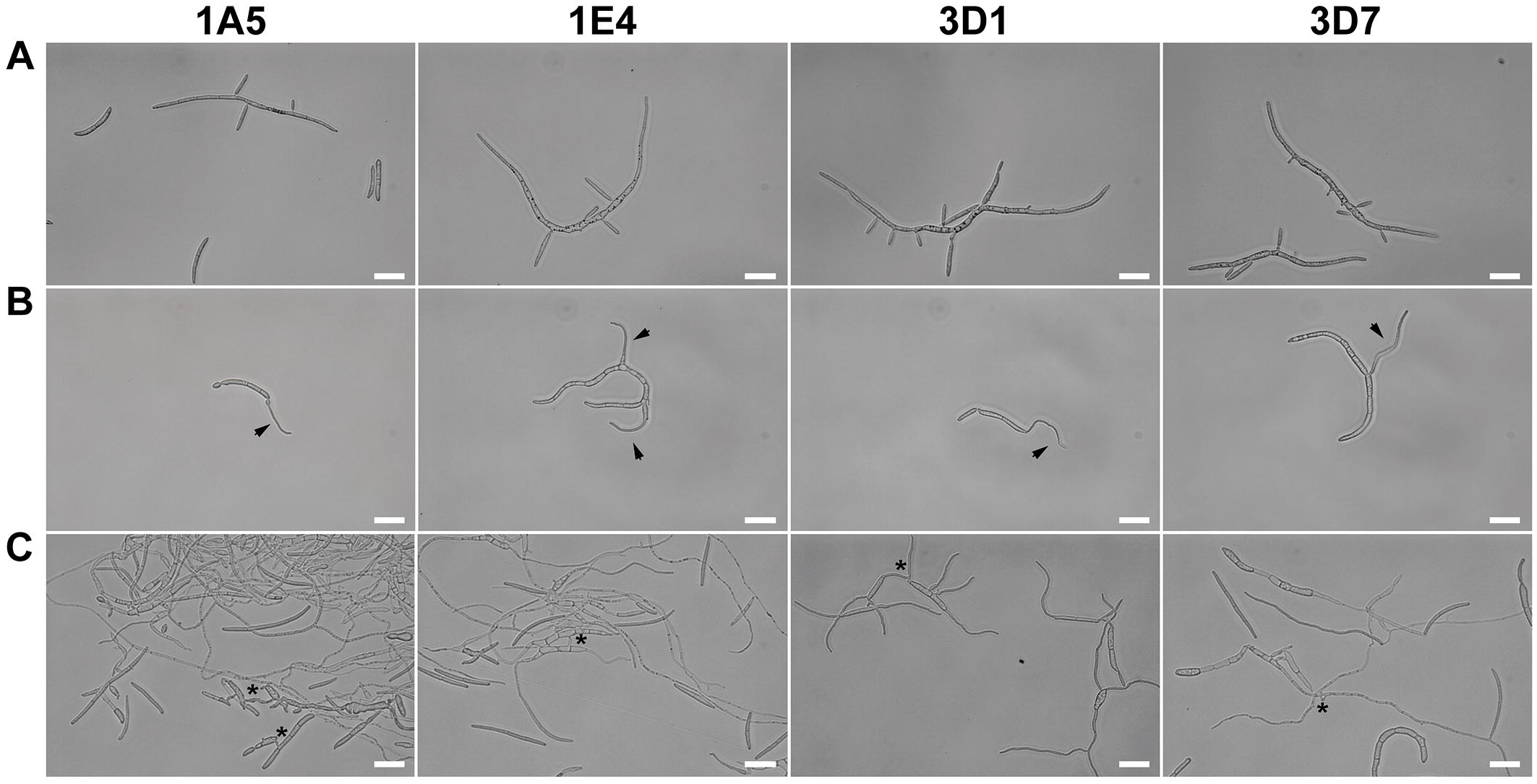
Effect of carbon limitation on the cell morphology of four *Zymoseptoria tritici* strains. A nutrient-rich medium (YSB) and a carbon-depleted medium (Minimal Media without sucrose) were used to monitor the growth form transitions. (A) Blastospores incubated on YSB continued to multiply as budding blastospores and were used as controls. (B) Under carbon limitation stress, the blastospores of all tested strains began growing as hyphae after 24 hours of incubation (arrows). (C) At 144 hai, a mycelial growth was predominant, with frequent vegetative cell fusion events (asterisks) among hyphae. Bars represent 20 μm.

In nitrogen-depleted media (MM without ammonium nitrate), three strains (1A5, 3D1, and 3D7) initially formed hyphae from one apical end, generating the first and second hyphal elongations within 72 hai (S1B Fig). 1E4 responded most quickly to a low nitrogen environment, with hyphae forming at 24 hai (S1A Fig) and a network of filamentous hyphae formed at 72 hai (S1B Fig). In regular MM all strains grew as a mixture of morphotypes (S1C Fig). Interestingly, the hyphae induced under both carbon and nitrogen limitation (Fig 2; S1 Fig) were thinner than those produced by high temperature (Fig 1).

### Oxidative stress induces morphological changes in some strains

To determine the effect of oxidative stress on the *Z. tritici* morphology we examined the growth in presence of hydrogen peroxide (H_2_O_2_) at two different spore concentrations. At a high initial cell density (10^5^ blastospores/mL), the presence of H_2_O_2_ stimulated hyphal growth in 1E4 up to a concentration of 5 mM (S2 Fig). 1A5, 3D1, and 3D7 continued to replicate as blastospores, but there was a substantial reduction in blastospore formation starting at 5 mM H_2_O_2_ (S2 Fig). To test whether a lower concentration of blastospores would change sensitivity to oxidative stress, we inoculated flasks with 10^3^ blastospores/mL. After 72 hours of incubation in 0.5 mM H_2_O_2_, strains 1E4 and 3D1 showed apical and lateral hyphal branching (S3 Fig), but both strains continued to form blastospores. At 1 mM, blastospores from 1E4, 3D1, and 3D7 developed into a mixture of pseudohyphae and regular hyphae (S3 Fig). The higher sensitivity at low cell densities suggests that the exogenous H_2_O_2_ was more rapidly degraded by secreted hydroperoxidase proteins (catalases, peroxidases and catalase-peroxidase) when cell densities were 100X higher. The 1A5 strain did not switch to a different morphotype at any H_2_O_2_ concentration, continuing to replicate as blastospores (S3 Fig). Starting at 2 mM H_2_O_2_, none of the strains changed their morphology (S3 Fig) and blastosporulation ceased (from 5 mM to 10 mM), suggesting a cytotoxic effect of H_2_O_2_ at these concentrations.

### Inoculum size effect suggests that quorum sensing may affect morphology

In order to evaluate the effect of the inoculum size on the morphological transition, we incubated all four strains at five cell densities in two growth conditions shown to induce hyphal growth. After 144 hours of incubation in C-depleted medium, we observed that morphology was independent of cell density for all four strains up to a concentration of 10^5^ blastospores/mL (S4A Fig), with most of the blastospores switching to hyphal growth and forming mycelial colonies. However, when the initial cell density was ≥10^6^ blastospores/mL, we observed a substantial reduction in the blastospore-to-hyphae transition and in hyphal length. No hyphae formed for 3D1 and 3D7 at initial cell densities of 10^7^ blastospores/mL, and they were clearly reduced for 1E4 and 1A5 (S4A Fig).

In YSB at 27°C we also found that initial inoculum size affected cell morphology (S4B Fig), though there were more pronounced differences among strains in this environment. At 144 hai, starting from the lowest initial cell density (10^3^ blastospores/mL), we observed pseudohyphae formation in 1A5, 1E4, and 3D1, and a mixture of pseudohyphae and chlamydospore-like cells in 3D7. At an initial density of 10^4^ blastospores/mL, the same phenotypes were found, except that 3D1 formed apical and lateral hyphae. Intermediate initial cell densities (10^5^ and 10^6^ blastospores/mL) promoted hyphal growth for 1E4, 3D1 and 3D7, but the 3D7 strain grew as a mixture of mycelium and chlamydospore-like cells at 10^6^ blastospores/mL. The increase of the inoculum size to 10^7^ blastospores/mL induced formation of chlamydospore-like structures in 1E4 and 3D1, and a transition from blastospores to pseudohyphae and chlamydospore-like cells in 3D7. 1A5 was not able to switch to hyphae under high temperatures in any of the initial cell densities (S4B Fig). The overall pattern of cell density dependence shown here suggests that a quorum sensing molecule may be controlling the morphological transition, as has been reported for other pleomorphic fungi [38–40].

### Mycelial growth is co-regulated with virulence-related genes

Aiming to identify genes associated with mycelial growth in nutrient-poor environments, we compared the transcriptomes from blastospores and two different hyphal inducing conditions for the four strains (S5Fig). The total number of sequencing reads and mapped reads per strain under different growth conditions is given in S1 Table. A total of 4888, 5296, 5323, and 6003 genes were differentially expressed among all tested conditions in 1A5, 1E4, 3D1, and 3D7, respectively. On average, 24% and 25% of the genes were upregulated during blastospore and mycelial growth, respectively.

The upregulated genes shared among all four strains for each morphology were selected based on pairwise comparisons. Two distinct sets of 134 genes shared among the four strains were identified in both comparisons. The first set is composed of genes that were upregulated during blastospore growth: these genes were upregulated in YSB compared to C-depleted medium, and were upregulated in YSB compared to 7dpi infection, FDR ≤0.01 (S6A Fig; S2 Table). The second set consists of genes that were upregulated during mycelial growth: these were upregulated in C-depleted medium compared to YSB, and were upregulated in wheat infection compared to YSB, FDR ≤0.01 (S6B Fig; S3 Table). For both comparisons, the sets of genes that do not overlap represent genes with a strain-specific expression profile.

Among the 134 shared genes that are significantly upregulated during blastospore growth, 20% are annotated as *hypothetical proteins* (Fig 3A; S2 Table). The remaining 107 genes are divided into fourteen biological process categories, with *metabolism* being the most represented term (44%). This category was composed mainly of non-secreted enzymes, including oxidoreductases (n=11), dehydrogenases (n=8), hydrolases (n=7), and transferases (n=9) (Fig 3A). No enrichment was found for any other category. However, seven genes from other categories were already described in other fungi to be involved in blastosporulation (S4 Table). As expected, genes related to vegetative blastospore growth were highly expressed in nutrient-rich medium - the environment most conducive for this morphotype (S6C Fig).

**Fig 3.**
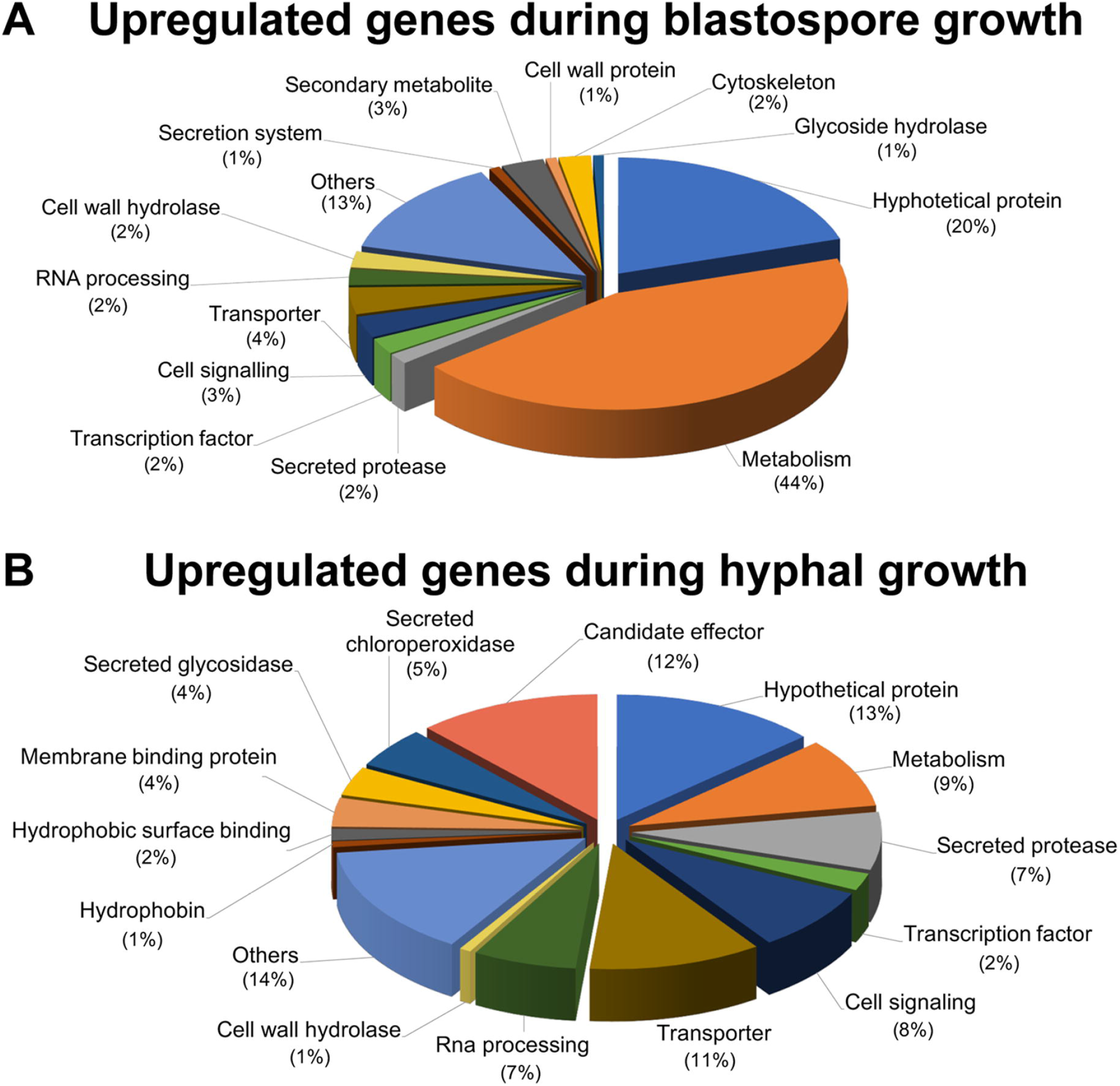
Transcriptome signatures of blastospore and mycelial growth phases organized based on enrichment of molecular functional terms. (A) Common upregulated genes during blastospore growth among the *Zymoseptoria tritici* strains. (B) Common upregulated genes for mycelial growth among the *Z. tritici* strains.

Among the 134 genes that were upregulated in all strains during mycelial growth, we observed an enrichment of secreted proteins (32%) known to be associated with fungal virulence, including *candidate effectors* (small secreted proteins - SSPs), *secreted proteases*, *secreted glycosidases*, *secreted chloroperoxidases*, *cell wall hydrolases*, *hydrophobins*, and *hydrophobic surface binding proteins* (Fig 3B; S3 Table). Overall, these virulence-associated genes were upregulated in C-depleted medium and during all stages of plant infection, albeit with a reduction in expression during the saprophytic phase of infection (at 28 dpi). The peak of expression of these virulence-associated genes preceded the first visual symptoms of infection, at 12 dpi for 1A5 and 3D7, and at 14 dpi for 1E4 and 3D1 (S6D Fig), as previously reported for some of these gene categories [41]. These patterns suggest that mycelial growth is co-regulated with the expression of genes relevant for virulence even in the absence of the host. We also detected the upregulation of six genes related to cellular morphogenesis (S5 Table). The transcription profiles of these genes showed an upregulation in both mycelial growth conditions (C-depleted medium and plant infection at 7 dpi) with the peak of expression occurring at 7 dpi for all four strains (S6E Fig).

### Chlamydospore-like cells are produced by *Z. tritici*

As mentioned above, we observed the formation of structures with the same morphology as chlamydospores. These structures had never been previously reported in *Z. tritici*. We aimed to determine whether these chlamydospore-like cells had properties associated with chlamydospores in other fungi. Chlamydospores are known to be survival structures that have thicker cell walls and a high content of lipid droplets [42]. We stained the different morphological cell types of 1A5 with the chitin-binding dye Calcofluor White (CFW) and the lipid droplets dye Nile Red (NR) to measure these properties. The intensity of CFW fluorescence was higher in chlamydospore-like cells (Fig 4A-4C) compared with blastospores, hyphae, and pseudohyphae that exhibited a weaker fluorescence for CFW (Fig 4). NR staining revealed that chlamydospore-like cells had a high content of lipid droplets, which was also observed for blastospores and pseudohyphae. On the contrary, hyphae showed a lower lipid droplet content than spores, with the exception of the hyphal branching zones (Fig 4) as has been reported for other fungi [43]. We then examined the thickness of the cell walls for three cell types using transmission electron microcopy (TEM). TEM showed that the chlamydospore-like cells had thicker cell walls than blastospores and pycnidiospores (Fig 5). We also found that pseudohyphae and the elongated suspensor cells, where chlamydospores are usually attached to the surrounding mycelium, have a cell wall thickness similar to pycnidiospores (Fig 5). No differences were observed for the chlamydospore-like cells produced by the 1A5 and 3D7 strains. On average, chlamydospore-like cells produced by *Z. tritici* had a cell wall thickness of ~460 nm, about four times thicker than the cell walls found in blastospores and pycnidiospores. Despite the thickness of the cell wall, we could not differentiate an extra cell wall layer as reported in chlamydospores from other fungal species [44,45].

**Fig 4.**
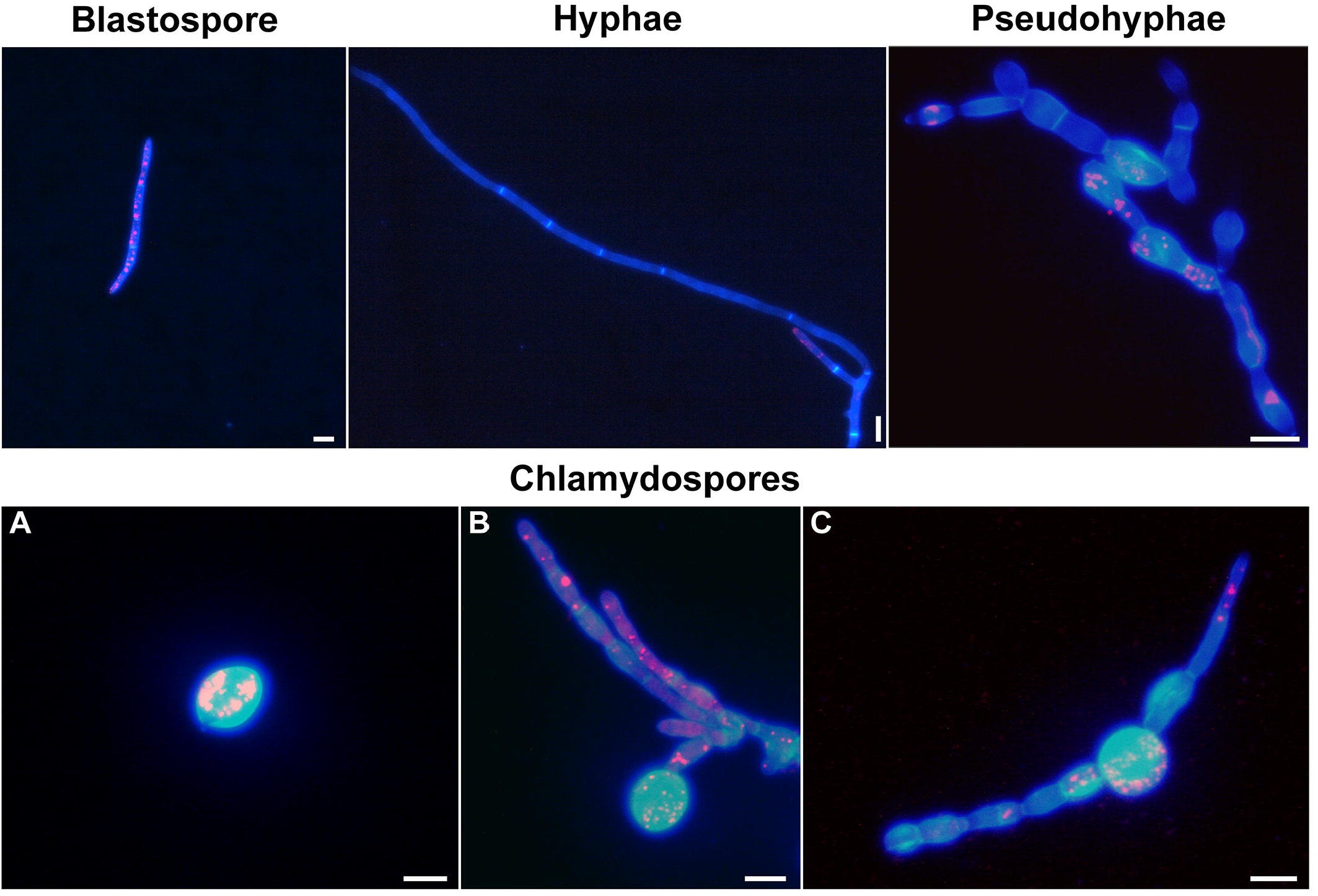
Staining of cell wall chitin and lipid content in the different morphotypes of *Zymoseptoria tritici*. Cell walls were stained with the chitin-binding dye Calcofluor White (blue color) and the lipid droplets were stained with Nile Red (red color). Blastospores, hyphae, and pseudohyphae showed a weaker fluorescence for Calcofluor White compared to chlamydospores (A), including those formed terminally (B) or intercalarily (C) via suspensor cells. A high accumulation of lipid droplets was observed in chlamydospores, as well as in blastospores and pseudohyphae. Hyphae only showed a high accumulation of lipid droplets in the hyphal branching zone. Bars represent 10 μm.

**Fig 5.**
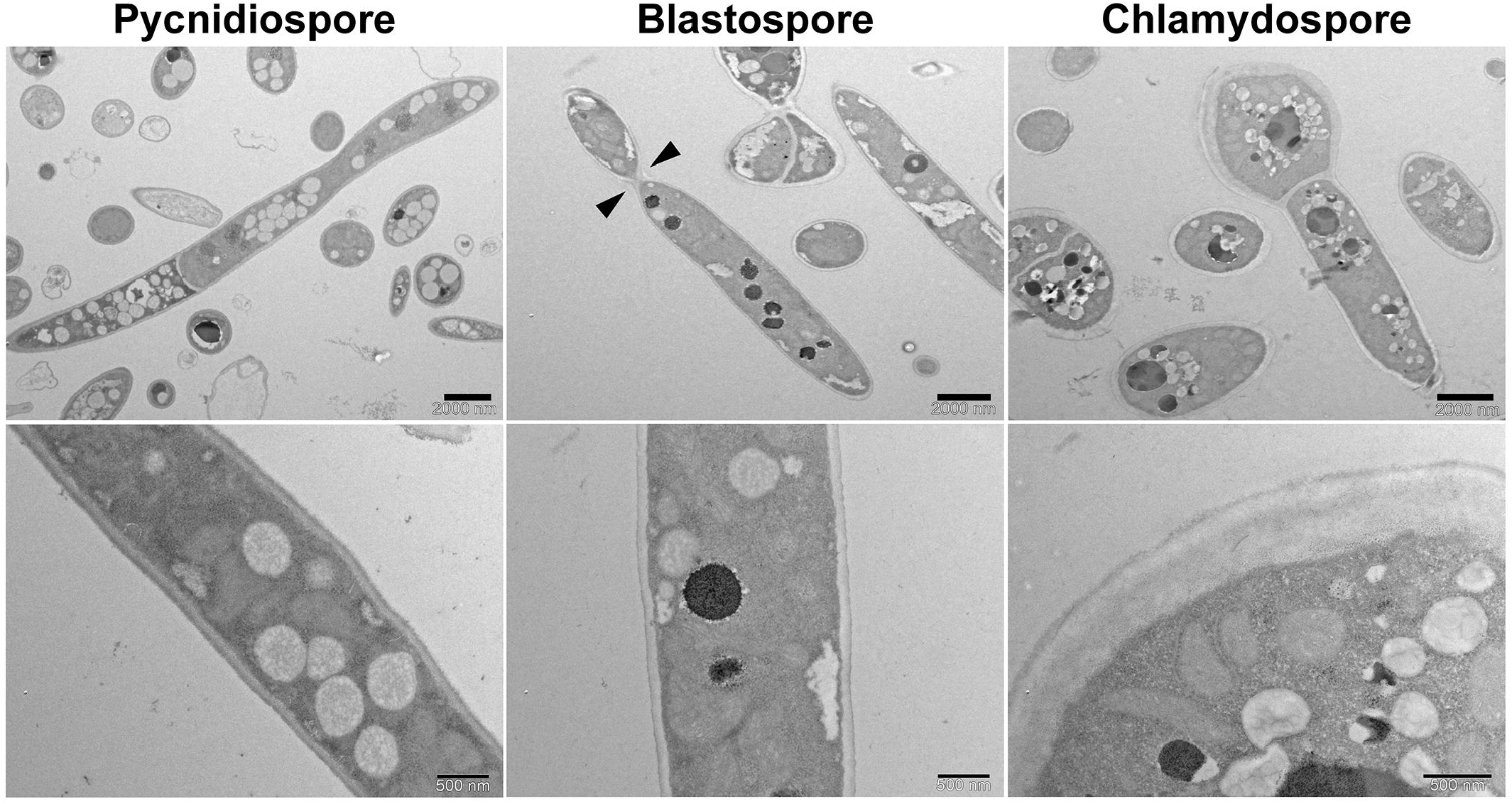
Chlamydospore cells produced by *Zymoseptoria tritici* have thicker cell walls than pycnidiospores and blastospores. Black triangles point to a detaching daughter cell budding from a blastospore mother cell. Images were obtained by transmission electron microscopy. Two different magnifications are shown for each morphotype: upper panel scale bar = 2000 nm (low magnification), lower panel scale bar = 500 nm (high magnification).

To determine if the chlamydospore-like cells are also produced during plant infection, wheat plants were inoculated with pycnidiospores of all four strains expressing cytoplasmic GFP and kept in a greenhouse chamber until pycnidia formation. Microscopic examination of the spore solution harvested from 20-day old infected leaves showed the presence of chlamydospore-like cells mixed among the pycnidiospores (S7 Fig).

### The chlamydospore-like cells are metabolically active and highly stress tolerant

The chlamydospore-like cells of 1A5 germinated and produced hyphae. After 12 hours of incubation on water agar (WA) plates, we observed germ tubes emerging from the chlamydospore-like cells, which led to the development of hyphae at 24 hai (Fig 6B and 6C). Blastospores used as controls initially germinated from one apical side, generating the first and second germ tubes within 12 hai (Fig 6A). We inoculated wheat with chlamydospore-like cells from GFP-tagged 1A5 to observe by confocal microscopy whether these cells could germinate on the leaf surface. At 24 hpi, free chlamydospore-like cells and chlamydospore-like cells still attached to a suspensor cell produced germ tubes and hyphae on the surface of wheat leaves (Fig 6D). These results suggest that chlamydospore-like cells may be able to cause infections.

**Fig 6.**
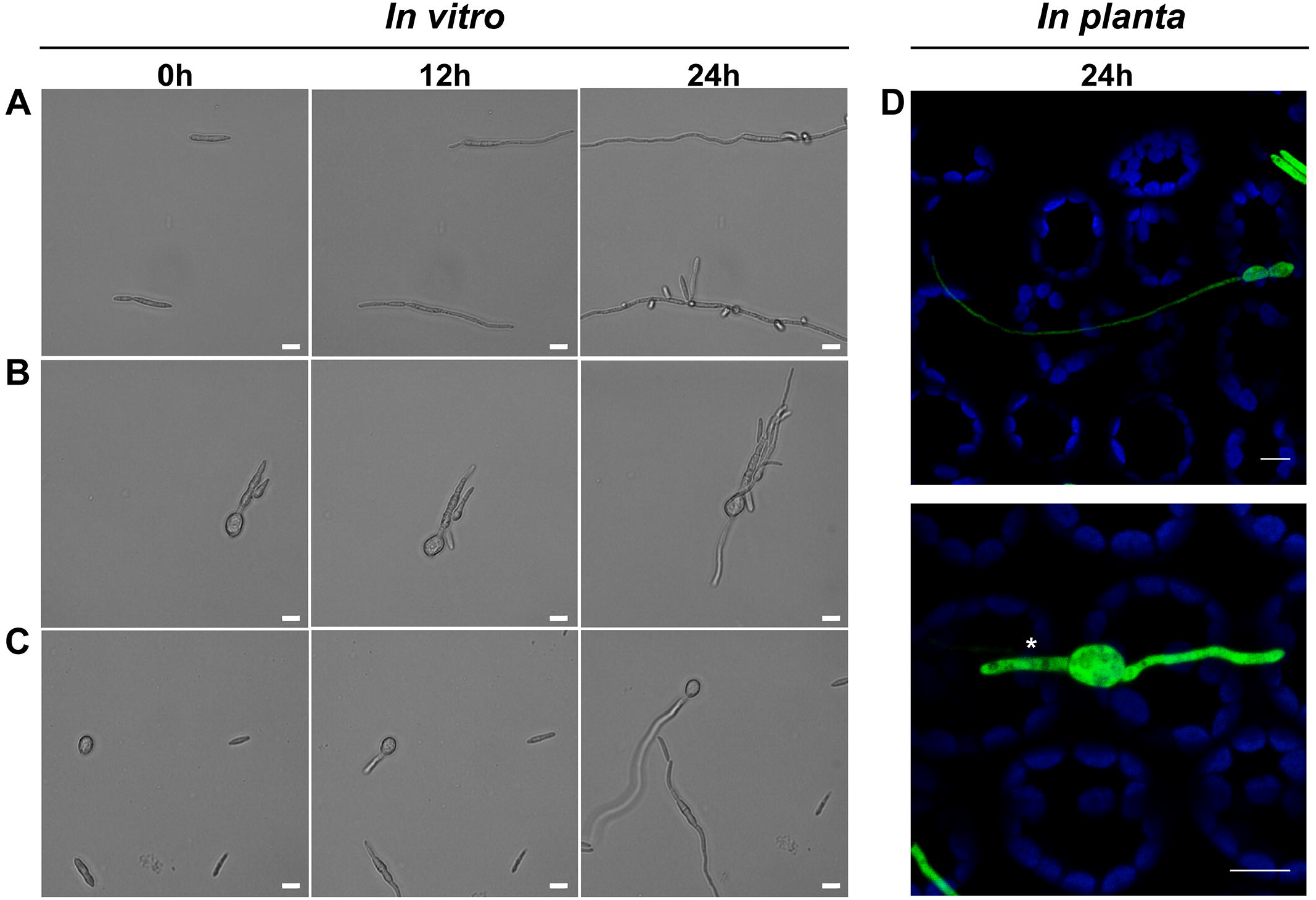
Chlamydospores produced by *Zymoseptoria tritici* are able to germinate both *in vitro* and on the surface of wheat leaves. (A) Blastospores were used as a control. Apical germination and hyphal elongation were observed after 12 hours of incubation on water agar (WA) plates at 18°C. At 24 hai, blastosporulation was observed from the germinated blastospores. (B) Terminal chlamydospores formed via suspensor cells and (C) free-chlamydospores began to germinate after 12 h of incubation on water agar. Hyphal elongation was obvious by 24 h. Bars represent 10 μm. (D) Free-chlamydospores (upper image) and terminal chlamydospores (lower image) expressing GFP (green color) produced germ tubes on the surface of wheat leaves at 24 hours after infection. Blue color corresponds to the autofluorescence detected from chlorophyll A. Asterisks indicate the suspensor cells where chlamydospores are attached. Bars represent 8 μm.

We also investigated whether chlamydospore-like cells share other characteristics of fungal resting spores, such as an ability to resist drought and temperature extremes. Incubation for one day under drought stress reduced viability of blastospores and pseudohyphae by 70%, while the chlamydospore-like cells had a reduction in cell viability of only 23% (Fig 7A). These rates changed little after 3 days of drought stress. After 5 days of drought stress, the survival rates were 10%, 19%, and 49% for blastospores, pseudohyphae and chlamydospore-like cells, respectively (Fig 7A). After 10 days of drought stress, the chlamydospore-like cells exhibited a survival rate of 50%, significantly higher than the other cell types (Fig 7A). In the cold stress experiment, only the chlamydospore-like cells survived the cold shock, with no statistically significant difference in the survival rates of 23%, 17%, and 19% after 1 min, 2 min and 3 min in liquid nitrogen, respectively (Fig 7B). In the heat stress experiment, 30°C affected blastospores and chlamydospore-like cells equally, but the 35°C treatment killed all the blastospores while chlamydospore-like cells and pseudohyphae survived at rates of 12% and 6%, respectively (Fig 7C). None of the morphotypes survived at 40°C (Fig 7C). For all tested environmental stresses, chlamydospore-like cells were the morphotype that exhibited the highest survival rate, suggesting that this structure exhibits ecological characteristics typical for fungal resting spores that enable improved survival to different stresses encountered in nature.

**Fig 7.**
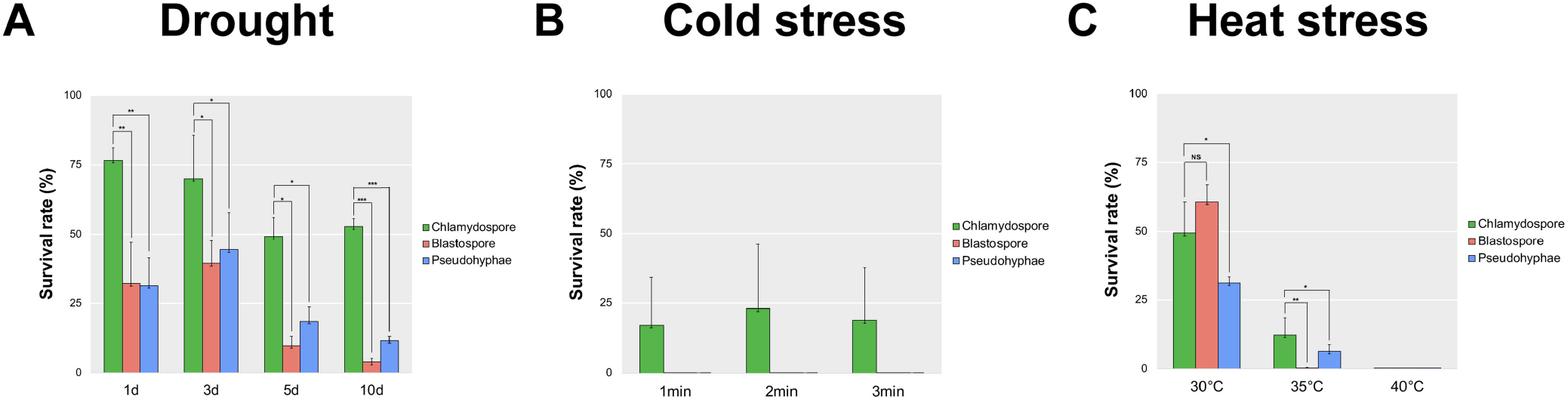
Chlamydospores produced by *Zymoseptoria tritici* are more tolerant to stress than blastospores and pseudohyphae. Survival rate (%) was estimated based upon the number of colonies formed after exposure to the stressful environment compared to the number of colonies formed in the control environment after seven days. (A) Blastospores, pseudohyphae and chlamydospore cells were kept for 1, 3, 5, and 10 days in a sealed box containing anhydrous silica gel to produce a dry environment. This simulated drought stress affected blastospore and pseudohyphae survival over time. Chlamydospores were less affected and exhibited a survival rate of 50% after 10 days under drought stress. (B) The same three morphotypes were subjected to a cold shock. Blastospores, pseudohyphae, and chlamydospores were submerged in liquid nitrogen for 1, 2, and 3 minutes at -196 °C. Only chlamydospores survived this cold stress. (C) The same three morphotypes were subjected to heat stress based on incubation for 24 hours at 30°C, 35°C or 40°C. Blastospores and chlamydospores were equally affected at 30°C. The incubation at 35°C killed all the blastospores, while chlamydospore survived at a 12% rate, double the rate observed for pseudohyphae (6%). No colonies were observed after incubation at 40°C. One star indicates that the adjusted p value is less than 0.05, two stars indicates that the p value is less than 0.005, and three stars indicates that the p value is less than 0.0005. ns indicates no significant difference.

### Germinating pycnidiospores produce blastospores on the surface of wheat leaves

Pycnidiospores of a GFP-tagged 3D7 strain were inoculated onto wheat plants and monitored using confocal microscopy. At 24 hai, we observed the expected pycnidiospore germination and initial hyphal branching, but we also observed newly-budding blastospores (Fig 8A). After 48 hours, there was an increase in the number of budding points on germinated pycnidiospores, and an increase in the number of free blastospores visible on the leaf (Fig 8B). This suggests that pycnidiospores of *Z. tritici* can germinate to produce additional blastospores even before penetration of the wheat leaf, providing an alternative mechanism for splash-dispersal of infective propagules and potentially providing a substantial increase in the total inoculum load associated with an infection cycle. At 48 hpi we also noticed many anastomosis events between pycnidiospores and new blastospores (Fig 8B).

**Fig 8.**
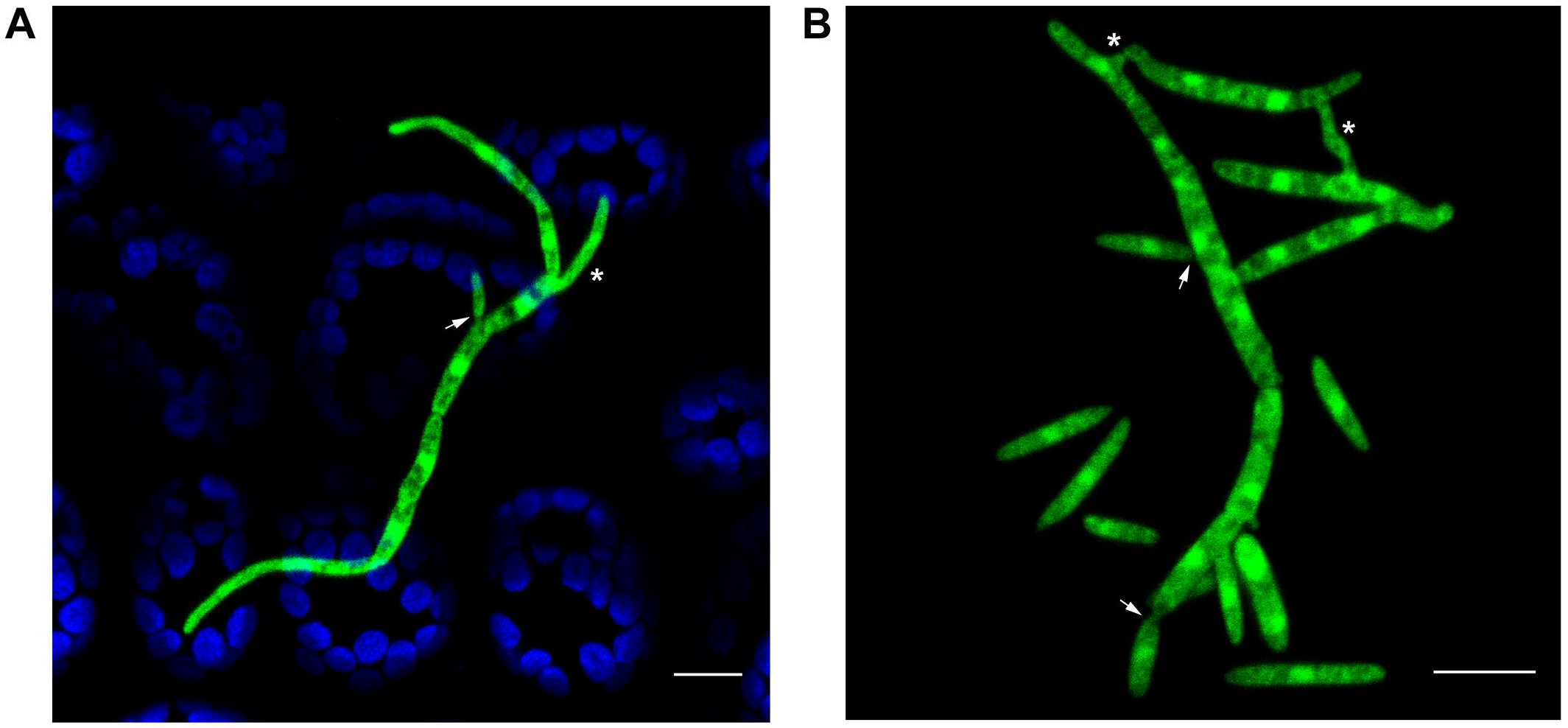
Germinated pycnidiospores of *Zymoseptoria tritici* produce new blastospores via blastosporulation on the surface of wheat leaves. (A) At 24 hours after infection, pycnidiospores (asterisk) germinated and formed hyphal branches expressing GFP (green color). The germinated pycnidiospores produced new blastospores on the host leaves (arrow). (B) We observed a significant increase in budding points on germinated pycnidiospores (arrows), as well as more free blastospores on the leaf surface at 48 hours after infection. At this time point, we also observed several vegetative cell fusion events between blastospores and pycnidiospores, which are characterized by a tubular shaped bridge between the fusing cells (asterisks). Blue color corresponds to autofluorescence detected from chlorophyll A. Bars represent 10 μm.

## Discussion

Fungi evolved to perceive a wide array of environmental signals and reprogram their cellular responses accordingly in order to maximize their fitness. Though much research has been oriented around morphological changes and their associated transcriptional pathways in response to different stressors in fungal pathogens of humans [10,15,46], knowledge in this area is limited for plant pathogens. Our study shows how different environmental stresses can affect cellular morphologies in the wheat pathogen *Z. tritici*. We also demonstrate the phenotypic plasticity of this fungus in response to environmental changes, as well as differential responses to the same environments among strains. An unexpected outcome of this study was the discovery of two novel morphotypes, one of which may have important epidemiological consequences.

We found that several environmental stresses can affect morphological transitions in strain-specific ways for four natural *Z. tritici* strains sampled from the same Swiss population. All four strains switched to hyphal growth in a nutrient-poor minimal medium. Morphological changes induced by limited nutrients have also been described in other dimorphic plant pathogenic fungi distributed across different phyla [9,47]. Nutrients are known to be present on the leaf surface in very small quantities and are usually surrounded by epicuticular waxes [48]. As a result, plant pathogens typically experience nutritient limitations while growing on leaves. It is thought that secreted enzymes, including plant cell wall degrading enzymes (PCWDEs) such as cutinases, are required to obtain nutrients during this stage of development, but *Z. tritici* has few PCWDEs compared to other fungi [49], suggesting that the perception of a nutrient-limited environment may act as a stimulus to induce a switch to an exploratory morphotype (hyphae) that can better explore the environment for food. We found a significant increase in the frequency of anastomosis events during mycelium formation under nutrient-limited conditions. Mycelial networks formed by anastomosis enable a shared cytoplasmic flow that can more efficiently distribute water, nutrients, and signals within a fungal colony [50,51]. Moreover, hyphal fusions were shown to be required for virulence for some plant pathogens [52,53]. We hypothesize that hyphal anastomosis provides benefits during epiphytic growth of germinating *Z. tritici* pycnidiospores prior to stomatal penetration, with anastomosis facilitating colony establishment and increasing strain fitness during colonization of the host.

The oxidative burst is a ubiquitous plant defense response to pathogen infection, resulting in a rapid release of reactive oxygen species (ROS), including hydrogen peroxide (H_2_O_2_) [54]. The ROS produced during plant defense were proposed to have various functions, including activation of plant defense-related genes and production of phytoalexins in addition to a direct antimicrobial effect that inhibits pathogen growth [55–57]. Though many studies have demonstrated its antimicrobial effects, the effects of H_2_O_2_ on morphological transitions remain largely unexplored. In *Candida albicans*, H_2_O_2_ induces hyphal differentiation in a dose-dependent manner [58]. Our results show that blastospores of three out of four *Z. tritici* strains switch to hyphae and pseudohyphae when exposed to H_2_O_2_ in a dose-dependent manner. A cell density dependence was also found. The different cell density responses suggest that the H_2_O_2_ is more rapidly metabolized by fungal hydroperoxidase proteins (catalases, peroxidases and catalase-peroxidase), leading to lower toxicity at higher cell densities. Several genes encoding peroxidases and catalase-peroxidases were identified in the genome of *Z. tritici* [59,60], which may explain the observed phenotype. The inability of 1A5 to differentiate hyphae from blastospores at any H_2_O_2_ concentration suggests that this strain either cannot sense oxidative stress or that it possesses a more efficient antioxidant response that can neutralize the effects of H_2_O_2_. Further experiments will be needed to test these hypotheses.

Temperature is an important abiotic factor that strongly affects growth, reproduction, and development in fungi [66,67]. Thermally dimorphic human fungal pathogens usually undergo morphological transitions in response to host temperature [47,68]. *Z. tritici* is the first plant pathogen shown to undergo morphological transitions after exposure to high temperatures [28,66]. Our experiments showed that a change in temperature from 18°C to 27°C triggered hyphal growth in three out of four tested *Z. tritici* strains. For these three strains, an increase in hyphal elongation and formation of branches occured soon after exposure to the higher temperature. The hyphae formed at a high temperature appeared thicker than those formed under nutritient limitations and it is not clear if they would be able to cause infections.

We used transcriptome sequencing to identify genes associated with mycelial growth in *Z. tritici*. It was shown in other plant pathogenic fungi that starvation induces the expression of virulence-related genes [69,70]. In *Z. tritici*, we found that genes relevant for virulence, including secreted hydrolytic proteins and candidate effectors, were upregulated in both hyphae-inducing conditions (carbon-depleted medium and *in planta*), suggesting that genes controlling morphogenesis and virulence may be coregulated under nutrient-limited conditions. In *C. albicans*, genes related to hyphal growth are co-regulated by transcription factors that affect expression of genes involved in virulence [5,7]. However, it is important to note that not every virulence function is connected to morphogenesis. The specific genes involved in the activation and regulation of these morphological transitions remain unknown for *Z. tritici*.

An unexpected outcome of this study was the discovery of chlamydospore-like cells in *Z. tritici*. The preponderance of evidence indicates that these cells are truly chlamydospores. Their morphology matched the chlamydospores described in other fungi: spherical, thick-walled cells with a high lipid content that form either intercalated or at the ends of filamentous hyphae and occasionally on pseudohyphae [17,71]. We also showed that these cells were able to survive several stresses that killed other cell types, including dessication and high and low temperatures, consistent with the function of chlamydospores as long-term survival structures in other fungi [16–18,72]. We conclude that *Z. tritici* produces chlamydospores.

This is the first time that chlamydospores have been found in a plant pathogen that is thought to mainly inhabit the phyllosphere. Though chlamydospores were especially prominent in the 1A5 strain at high temperature, we observed chlamydospore production in all four strains both *in vitro* and *in planta*. We hypothesize that the chlamydospores produced by *Z. tritici* may contribute an important source of inoculum that can survive between growing seasons and perhaps form a long-term spore-bank that can persist for many years in wheat-growing regions. Chlamydospores may also contribute to long-distance movement of the pathogen on infested straw or seeds. These properties could impact the management of septoria tritici blotch (STB), but additional studies will be needed to understand the roles played by chlamydospores during the STB disease cycle.

Another finding with epidemiological implications was to demonstrate that blastospores can form from germinating pycnidiospores on the surface of wheat leaves. This possibility was proposed more than 30 years ago [73,74], but was not followed up in subsequent studies. The formation of blastospores during *in vitro* growth in nutrient-rich medium is why *Z. tritici* has long been considered to be a dimorphic fungus, but until now it was not clear if the blastospores could also form on leaf surfaces in nature. Our experiments confirmed that blastospores are produced during a natural infection. It remains unclear whether blastospores make a significant contribution to the development of epidemics. Given their small size and increasing abundance over time, we hypothesize that they may play an important role in the explosive increase in STB that can occur following prolonged rainy periods that last for several days [73,75].

In conclusion, we demonstrate that a fungal plant pathogen responds morphologically to diverse environmental stresses, and we propose distinct biological roles for different morphotypes. We also illustrate the advantage of including different natural strains when studying the response of a species to environmental stresses, as we found different morphological responses among the tested strains. Our findings provide important new insights into the *Z. tritici*-wheat pathosystem including: 1) the phenotypic plasticity of *Z. tritici* in response to different environmental changes; 2) the reclassification of *Z. tritici* as a pleomorphic fungus due to its ability to produce four distinct morphotypes; 3) the discovery of chlamydospores that may help the fungus to survive stressful conditions; 4) the formation of blastospores from germinating pycnidiospores on the surface of wheat leaves, enabling a potentially significant increase in inoculum size. These findings position *Z. tritici* as an excellent model organism to explore how morphological changes can affect adaptation to different environments in fungal plant pathogens.

## Materials and methods

### Fungal strains, environmental stimuli, and growth conditions

Four previously characterized *Zymoseptoria tritici* strains (ST99CH_1A5, ST99CH_1E4, ST99CH_3D1, and ST99CH_3D7, abbreviated as 1A5, 1E4, 3D1, and 3D7, respectively); [76,77], were used in this study. Strains expressing cytoplasmic GFP [78] were provided by Andrea Sanchez-Vallet after being generated by Sreedhar Kilaru and Gero Steinberg as described in Zala *et al* 2018 [79]. All strains were grown in YSB (yeast extract 10 g/L and sucrose 10 g/L; pH 6.8). Depending on the tested environment, we used either YSB or a defined salts medium (Minimal Medium – MM, pH 5.8; [80]. Each strain was stored in glycerol at -80°C until required and then recovered in YSB medium incubated at 18°C for three days. For nutrient limitation experiments, the cells were washed three times with sterile distilled water to remove any remaining nutrients prior to placing them into nutrient-limited conditions. Cell concentrations were determined by counting blastospores using a KOVA cell chamber system (KOVA International Inc., USA).

### Effect of temperature on growth morphology

Blastospores of each strain were inoculated on YSB at a final concentration of 10^5^ blastospores/mL. Three flasks were incubated at 18°C (as control) and three at 27°C for four days. An aliquot was taken from each flask at 8, 24, 48, 96, and 144 hai to monitor cell morphology by light microscopy using a Leica DM2500 microscope with LAS version 4.6.0 software. These experiments were repeated three times.

### Effect of nutrient limitation on growth morphology

MM was used to monitor the transition in growth morphologies under different nutrient limitations. The effect of carbon depletion was assessed by adding each strain at a final concentration of 10^5^ blastospores/mL into either regular MM, which contains sucrose as the main carbon source, or into MM without sucrose (C-depleted medium) and incubated at 18°C. The cell morphology was observed by light microcopy every 24 hours until 144 hai. The effect of nitrogen depletion was determined using MM lacking ammonium nitrate, which is normally present in MM as the main nitrogen source. We compared the growth morphologies of cells growing in MM with and without ammonium nitrate by light microcopy during 96 hours of incubation at 18°C. The growth morphologies in both nutrient-limited conditions were compared to the morphologies found in the nutrient-rich YSB. Experiments were repeated three times for each nutrient environment.

### Effect of oxidative stress on growth morphology

H_2_O_2_ was added into YSB at different concentrations (0, 0.5, 1, 2, 5, and 10 mM) to induce different levels of exogenous oxidative stress. Each strain was added to a final concentration of 10^5^ blastospores/mL and incubated at 18°C. Morphological changes and cellular morphologies were observed by light microscopy at 72 hai. To determine whether a lower concentration of blastospores promotes a higher susceptibility to oxidative stress, we performed the same experiment using a final concentration of 10^3^ blastospores/mL. These experiments were repeated twice.

### Effect of inoculum size on growth morphology

Flasks containing YSB or C-depleted medium were inoculated with a final cell density of 10^7^ blastospores/mL, followed by a 10X serial dilution to a concentration of 10^3^ blastospores/mL. Flasks containing C-depleted medium were incubated at 18°C and flasks with YSB were incubated at 27°C. Cell morphology was determined every 24 hours using light microscopy over 144 hours of incubation. These experiments were repeated three times.

### RNA extraction, library preparation, and sequencing

Flasks containing a final concentration of 10^5^ blastospores/mL in YSB or C-depleted medium were incubated at 18°C to induce blastosporulation or mycelial growth, respectively. Total RNA was extracted by using TRIzol (Thermo Fisher Scientific, Waltham, Massachusetts) following the manufacturer’s instructions. RNA quality was checked using a Qubit fluorometer (Thermo Fisher Scientific, Waltham, Massachusetts) and a TapeStation 2200 (Agilent, Santa Clara, California). Truseq stranded mRNA kit (Illumina Inc., San Diego, CA, USA) was used for library preparation, and ribosomal RNA was depleted by polyA enrichment. The quality and quantity of the enriched libraries were assessed using a Qubit fluorometer (Thermo Fisher Scientific) and TapeStation (Agilent). Libraries were sequenced on the Illumina HiSeq 2500 at 4 x 100 bp paired end reads.

### Transcriptome mapping and quantification

The sequencing adapters and reads shorter than 15 bp were trimmed from raw Illumina reads using Trimmomatic v.0.36 [81] with the following settings: ILLUMINACLIP:TrueSeq3-PE.fa:2:30:10, LEADING:2 TRAILING:2, SLIDINGWINDOW:4:15 MINLEN:15. Filtered reads of each *Z. tritici* strain for the most common growth morphologies (blastospores and mycelial growth) were mapped against their respective genomes using Tophat v.2.0.13 [82]. HTSeq-count v.0.6.1 was used to calculate the gene counts [83]. Mapped reads with more than one reported alignment were excluded from further analyses. We applied the TMM (trimmed mean of M values) method implemented in the Bioconductor edgeR v.3.12.0 package to normalize the gene counts and to calculate the differentially expressed genes under each condition [84,85]. Only genes with average counts per million reads per sample >1 for at least one growth condition were counted as expressed genes. The Benjamin-Hochberg false discovery rate (FDR) correction was used to adjust p-values based on the Fisher exact test.

### Transcription profile analysis

We generated the CPM (counts per million mapped reads) values based on TMM-normalized library sizes obtained from the edgeR v.3.12.0 package [84]. Only genes with adjusted p-values of FDR≤0.01 were included in the analysis. The published transcriptomes of each strain during wheat infection at 7 dpi [41] were used for comparative analyses (datasets in NCBI SRA SRP077418). Only genes that were significantly upregulated during both wheat infection at 7 dpi and in C-depleted medium compared to YSB were considered to be mycelial growth-related genes. These genes were used in further analyses. Specific transcriptome signatures were identified based on the differentially expressed genes that were shared among all four strains under each growth condition, according to the *Z. tritici* reference genome annotations [86,87]. We also used the published transcriptomes of different stages of wheat infection [41] to interpret the expression profiles of some genes under both axenic and *in planta* conditions. The log_2_-transformed CPM values of those genes were plotted using the R package ggplot2 [88].

### Fluorescence microscopy to quantify chitin and lipids in different morphotypes

Blastospores of 1A5 were inoculated into YSB to a final concentration of 10^5^ blastospores/mL and incubated at 18°C or 27°C to induce blastosporulation or chlamydospore-like formation, respectively. 1A5 was also inoculated at the same concentration into C-depleted medium and incubated at 18°C to induce hyphal growth. After four days of incubation, the resulting cells were harvested and fixed with 70% (v/v) ethanol for 30 minutes, followed by three washes with phosphate buffer saline (PBS). The cells were stained with 1 μg/mL of CFW and 2.5 μg/mL of NR (both reagents from Sigma-Aldrich Chemie Gmbh, Munich, Germany) for 15 minutes. The stained cells were viewed with a Leica DM2500 fluorescence microscope with LAS version 4.6.0 software, using a UV filter system for CFW consisting of a BP excitation filter at 340-380 nm and a long pass emission filter (>425 nm), and a red filter system for NR consisting of a BP excitation filter at 546/10 nm and an emission filter at 585/40 nm.

### Transmission electron microscopy

Cells of 1A5 and 3D7 were collected after 4 days of growth in YSB at 18°C or 27°C. Pycnidiospores for each strain were harvested from infected wheat plants. Samples were prepared using microwave-assisted fixation in 0.1 M cacodylate buffer (2.5% glutaraldehyde and 2% paraformaldehyde), using a PELCO Biowave (Ted Pella, California, USA), followed by embedding in 2% low melting temperature agarose (SeaPlaque Agarose, FMC Corporation, USA). A second step of microwave-assisted fixation was used after embedding in agarose. Before resin infiltration, the samples went through two steps of post-fixation, first in osmium tetroxide (OsO4) in 0.1 M cacodylate buffer and then in 1% uranyl acetate (UrAc), followed by a graded series of ethanol washes for dehydration (25%, 50%, 75%, 90%, and 100%). Samples were infiltrated with Spurr resin and polymerized at 60°C for 48 hours. Ultrathin sections (60 nm) were cut using the Leica Ultracut FC6 (Leica Microsystem, Vienna) and then transferred to formvar coated carbon grids, followed by post-staining with uranyl acetate and lead citrate. Micrographs were collected using a Morgagni 268 transmission electron microscope operating at 100 kV and using a CCD (1376×1032 pixel) camera.

### Germination of chlamydospore-like cells and their survival under stress

Chlamydospore-like cells and blastospores of the 1A5 strain were harvested from YSB after 4 days of growth at 27°C or 18°C, respectively, and their concentrations were adjusted to ~10^5^ cells/mL. 10 μL was spotted onto WA (1% agar in water) and the plates were incubated at 18°C. Germ tube formation was checked at 0, 12, and 24 hours after inoculation by light microscopy.

To determine whether chlamydospore-like cells differ in their ability to survive environmental stresses compared to blastospores and pseudohyphae, we exposed 8×10^3^ cells/mL of each morphotype to different stresses, including drought and extreme temperatures.

For drought stress, 200 μL of each cell solution were spread onto Amersham Hybond™ N^+^ nylon membranes (GE Healthcare Life Sciences, Chicago, Illinois) and dried in a sterile Petri dish placed on a sterile bench in a laminar flow hood. The Petri dishes were then placed into a sealed box containing anhydrous silica gel for 1, 3, 5, or 10 days. After the drought stress, the membranes were placed onto YMA plates (4g/L yeast extract, 4g/L malt extract, 4g/L sucrose, and 12g/L agar), incubated at 18°C and colony formation was monitored. Control membranes containing each morphotype were transferred directly to YMA plates without any drying treatments.

For cold stress, 300 μL aliquots of each cell solution were submerged into liquid nitrogen for 1, 2, or 3 minutes, and 200 μL of the thawed cell solution were spread onto YMA plates. Control samples were incubated at room temperature. For heat stress, 200 μL aliquots of each morphotype were incubated for 24 hours at 30, 35, or 40°C, followed by plating on YMA. Controls were kept at 18°C. YMA plates from control and treated samples were incubated at 18°C for seven days. Survival rates for each cell type were estimated based on the number of colonies formed after exposure to the stressful conditions compared to the number of colonies formed by the controls.

### Plant infection assays

The susceptible wheat cultivar Drifter was used in all experiments. Plant growth conditions were described earlier [89]. All infection assays used 16-day-old plants. For all four strains, blastospore suspensions produced after 3 days of growth in YSB at 18°C were filtered through two layers of cheesecloth and diluted to a final concentration of 10^6^ blastospores/mL in 30 mL of sterile water supplemented with 0.1% (v/v) Tween 20. Spore suspensions were applied to run-off using a sprayer and the plants were kept for three days in sealed plastic bags, followed by 21 days in a greenhouse. Leaves with pycnidia were harvested and transferred to a 50 mL Falcon tube containing sterile water and gently agitated to harvest the pycnidiospores. Pycnidiospore suspensions were adjusted to a final concentration of 10^6^ pycnidiospores/mL and a new batch of plants were inoculated as described above. The plants infected by pycnidiospores were used to observe the formation of chlamydospore-like cells on wheat plants.

To determine whether chlamydospore-like cells produce germ tubes and initiate hyphal growth after coming into contact with the host, we inoculated Drifter with chlamydospore-like cells from the 1A5 strain tagged with cytoplasmic GFP. Production of chlamydospore-like cells was induced using high temperature (YSB at 27°C) as described above. The suspension of chlamydospore-like cells was prepared and concentrations were standardized as described for blastospore infections. The inoculation procedure was the same except the inoculated plants were kept in plastic bags for only 24 hours.

To observe formation of chlamydospore-like cells *in planta*, infected leaves from wheat plants inoculated with pycnidiospores were harvested at 20 dpi. The infected leaves were incubated in a humidity chamber for 24 hours, following by washes with sterile distilled water to collect the released spores. The spore suspension was checked for the presence of chlamydospore-like cells using the fluorescent microcope with a GFP filter consisting of a BP excitation filter at 480/40 nm and a BP emission filter at 527/30 nm.

To observe blastospore formation on the surface of wheat leaves, we inoculated plants using blastospores of the 3D7 strain tagged with cytoplasmic GFP. At 21 dpi, GFP-tagged 3D7 pycnidiospores were harvested and a new batch of plants was inoculated using these pycnidiospores as described earlier. The inoculated plants were kept in plastic bags at 100% humidity until they were examined for blastospores at 24 and 48 hai.

### Confocal microscopy of germinated pycnidiospores and chlamydospore-like cells

Confocal images were obtained using a Zeiss LSM 780 inverted microscope with ZEN Black 2012 software. An argon laser at 500 nm was used to excite GFP fluorescence and chloroplast autofluorescence with an emission wavelength of 490-535 nm and 624-682 nm, respectively. Plants inoculated with GFP-tagged 1A5 chlamydospore-like cells were checked at 24 hai to observe germination of the chlamydospore-like cells. The same confocal settings were used to determine whether pycnidiospores produced blastospores on the leaf surface. Wheat plants inoculated with GFP-tagged 3D7 pycnidiospores were checked at 24 and 48 hai.

## Acknowledgements

We acknowledge Gero Steinberg and Sreedhar Kilaru for generating the fluorescent *Z. tritici* Swiss strains used in this study, and Andrea Sanchez-Vallet for providing the strains. We thank Daniel Croll and Andrea Sanchez-Vallet for critical reading of the manuscript. We also thank Andrea Sanchez-Vallet, Lukas Meile, Julien P. L. Alassimone, and Danilo A. Pereira for discussions in the course of this work. RNA processing and sequencing were performed in the Genetic Diversity Center (GDC) of ETH Zurich and the Genomics Facility of ETH Basel, respectively. Transmission electron microscopy (TEM) and confocal microscopy were performed at the Scientific Center for Optical and Electron Microscopy (ScopeM) of ETH Zurich.

## Supporting information

**S1 Fig. Effect of nitrogen depletion stress on the cell morphology of four *Zymoseptoria tritici* strains.** A nutrient-poor medium (MM without ammonium nitrate) was used to monitor the growth form transition. (A) At 24 hai, only blastospores from the 1E4 strain switched to hyphal growth forming lateral filaments (black arrow). The rest of the strains remain undifferentiated. Bars represent 10 μm. (B) At 72 hai, blastospores of all tested *Z. tritici* strains incubated under nitrogen depletion undergo a blastospore-to-hyphae transition. Branching and hyphal elongation were predominantly observed at this time point. Bars represent 30 μm. (C) Regular MM containing ammonium nitrate induced a mixture of morphotypes, such as blastospores, hyphae and budding blastospores from filamentous hyphae. Bars represent 30 μm.

**S2 Fig. Effect of exogenous oxidative stress induced by H_2_O_2_ on the cell morphology of four *Zymoseptoria tritici* strains using a high blastospore inoculum density (10^5^ blastospore/mL).** A nutrient-rich medium (YSB) containing different concentrations of H_2_O_2_ was used to incubate the strains under oxidative stress for 72 hours. Blastopores kept on YSB medium without addition of H_2_O_2_ continue to multiply as budding blastospores via blastosporulation and were used as a control (A, B, C, D - bars represent 50 μm). Despite the increases in H_2_O_2_ concentration in the medium, no morphological changes were observed for 1A5 (E, I, M, Q – bars represent 50 μm), 3D1 (G, K – bars represent 50 μm; O and S – bars represent 10 μm), and 3D7 (H, L, P, T - bars represent 50 μm). The 1E4 strain responded morphologically to H_2_O_2_ concentrations at 0.5 mM, 1 mM, and 5 mM (F, J, N - bars represent 50 μm) undergoing the blastospore-to-hyphae transition. As observed for other strains (Q, S, and T), blastospores of the 1E4 strain remained as undifferentiated structures at 10 mM (R - bars represent 50 μm).

**S3 Fig. Effect of exogenous oxidative stress induced by H_2_O_2_ on the cell morphology of four *Zymoseptoria tritici* strains using a low blastospore inoculum density (10^3^ blastospore/mL).** A nutrient-rich medium (YSB) containing different concentrations of H_2_O_2_ was used to incubate the strains under oxidative stress for 96 hours. Blastopores kept on YSB medium without addition of H_2_O_2_ continued to multiply as budding blastospores via blastosporulation and were used as a control. At 0.5 mM, only the 1E4 and 3D1 strains showed hyphal branching and the initial formation of mycelium. At 1 mM, blastospores from 1E4, 3D1, and 3D7 grew as a mixture of pseudohyphae (asterisks) and filamentous hyphae (black arrows). At 2 mM, none of the strains changed their morphology nor were they able to asexually reproduce via blastosporulation. Blastospores of the 1A5 strain did not switch to different morphotypes in any of the tested concentrations of H_2_O_2_. Bars represent 10 μm.

**S4 Fig. Cell morphology of four *Zymoseptoria tritici* strains after 144 hours of incubation on two distinct hyphal induction media and at different initial cell densities.** (A) Blastospores incubated in carbon-depleted MM at 18°C demonstrated a morphological independence of the inoculum size until the concentration of 10^5^ blastospores/mL, where they were able to switch to hyphal growth and to form mycelial colonies. At higher initial inoculum concentrations (≥10^6^ blastospores/mL), a drastic reduction of growth form transition and in hyphal length was observed for all tested strains. (B) Blastospores incubated in YSB medium at 27°C showed significant differences among the strains. At lower initial inoculum concentrations (10^3^ and 10^4^ blastospores/mL) only pseudohyphae and chlamydospores formed. Intermediate inoculum concentrations (10^5^ blastospores/mL and 10^6^ blastospores/mL) promoted hyphal growth for the 1E4, 3D1 and 3D7 strains. A ten-fold increase in the initial cell density (10^7^ blastospores/mL) favored chlamydospore formation in all strains. 1A5 blastospores were not able to switch to hyphae and undergo transitions to pseudohyphae and chlamydospores for all tested concentrations. Bars represent 10 μm.

**S5 Fig. Morphological forms of four *Zymoseptoria tritici* strains grown in two distinct morphotype-induction conditions used for the RNA-Seq.** Nutrient-rich medium (YSB) induces only blastosporulation. Nutrient-poor medium (MM) and the wheat leaf surface induced the blastospore-to-hyphae transition and mycelial growth in all tested strains. Wheat leaves infected with *Z. tritici* strains expressing cytoplasmic GFP (green color) were analyzed using confocal microscopy. The blue color corresponds to chlorophyll A detected from chloroplast autofluorescence. Bars represent 10 μm.

**S6 Fig. Core transcriptome signatures of four *Zymoseptoria tritici* strains from two distinct growth phases.** (A-B) Venn diagram of the overexpressed genes (FDR≤0.01) for each strain grown under blastospore and mycelial growth conditions. (C) Graphic representing average expression (log2-transformed counts per million values) of seven genes related to blastosporulation during the blastospore growth phase among strains. (D) Graphic representing average expression (log2-transformed counts per million values) of 38 virulence-related genes during the mycelial growth phase. (E) Graphic representing average expression (log2-transformed counts per million values) of six genes related to cellular morphogenesis during the mycelial growth phase.

**S7 Fig. Chlamydospore cells observed in a spore solution harvested from 20-day-old infected leaves.** Plants were previously inoculated with pycnidiospores of four *Z. tritici* strains. Free chlamydospores expressing cytoplasmic GFP were frequently observed mixed with pycnidiospores in the spore solution of all tested strains. Bars represent 30 μm. The proportion of chlamydospores was visibly lower compared to pycnidiospores.

**S1 Table.** Overview of the transcriptomic data of four *Zymoseptoria tritici* strains obtained from two distinct growth phases (blastospore *vs*. mycelium).

**S2 Table.** The shared 134 genes overexpressed during blastospore growth phase among four *Zymoseptoria tritici* strains.

**S3 Table.** The shared 134 genes overexpressed during mycelial growth phase among four *Zymoseptoria tritici* strains.

**S4 Table.** Expression values of seven genes overexpressed in blastospore growth phase and associated with blastosporulation.

**S5 Table.** Expression values of 39 genes overexpressed in mycelium growth phase and associated with virulence factors.

**S6 Table.** Expression values of six genes overexpressed in mycelium growth phase and associated with cellular morphogenesis.

## Accession numbers

All raw sequencing data were deposited into the NCBI Sequence Read Archive under the accession number SRP152081.

## Author contributions

**Conceptualization:** Carolina Sardinha Francisco, Javier Palma-Guerrero

**Formal analysis:** Carolina Sardinha Francisco, Xin Ma, Javier Palma-Guerrero.

**Funding acquisition:** Bruce A. McDonald

**Investigation:** Carolina Sardinha Francisco, Maria Manuela Zwyssig, Javier Palma-Guerrero.

**Methodology:** Carolina Sardinha Francisco, Javier Palma-Guerrero.

**Project administration:** Javier Palma-Guerrero

**Resources:** Bruce A. McDonald

**Supervision:** Javier Palma-Guerrero, Bruce A. McDonald

**Validation:** Carolina Sardinha Francisco

**Writing – original draft:** Carolina Sardinha Francisco, Javier Palma-Guerrero, Bruce A. McDonald.

**Writing – review & editing:** Carolina Sardinha Francisco, Javier Palma-Guerrero, Bruce A. McDonald.

